# GSLab: Open-Source Platform for Advanced Phasor Analysis in Fluorescence Microscopy

**DOI:** 10.1101/2025.02.10.637545

**Authors:** Alexander Vallmitjana, Belén Torrado, Amanda F. Durkin, Alexander Dvornikov, Navid Rajil, Suman Ranjit, Mihaela Balu

## Abstract

GSLab addresses the need for effective image analysis tools in fluorescence microscopy by providing an open-source platform that enhances traditional phasor analysis with advanced features. Key capabilities include machine learning-based clustering, real-time monitoring, and quantitative unmixing of fluorescent species. Designed for both commercial and custom systems, GSLab provides researchers with comprehensive lifetime and spectral phasor image analysis tools to tackle complex biological problems.

## Main Text

Fluorescence lifetime microscopy (FLIM) is a powerful imaging technique that provides quantitative information about the local environment and molecular interactions of fluorophores within biological tissues or other materials. By measuring how long a fluorescent molecule remains in an excited state before emitting a photon, FLIM offers a new dimension for biological imaging, which, combined with spectral imaging, enables researchers to investigate complex cellular processes and interactions. The integration of advanced analytical methods, such as the phasor approach, enables more sophisticated image analysis and interpretation. As a result, this will further enhance the capabilities of FLIM and spectral microscopy.

The basis of phasor analysis lies in using the phasor transform to map every pixel of an image onto a two-dimensional space known as phasor space, based on the photon distribution within that pixel across the fluorescence lifetime or spectral dimensions^1,2^. The position of each pixel in phasor space is determined by the shape of the photon distribution and is independent of the signal’s intensity. Analysis by means of the phasor representation does not require prior knowledge of the nature of the sample nor fitting of a model. In addition, utilization of the Fast Fourier Transform algorithm enables rapid computation. This analysis simplifies visual inspection and identification of distinct populations of pixels, which can subsequently be mapped back to the original fluorescence image (or set of images)^3^. Furthermore, the mathematical properties of the phasor transform enable researchers to understand the phenomena occurring in the sample by observing changes in the photon distribution represented in phasor space. A brief overview of the mathematics behind the phasor approach for analyzing fluorescence lifetime microscopy images is available in Online Methods.

Phasor analysis is widely used in the biophysics and bioimaging fields and has led to a vast number of quantification methods and applications. These include studying cellular metabolic states^4^, molecular interactions^5^, molecular dynamics^6^, drug delivery^7^, chromatin compaction^8^, sensing local polarity^9^, ion concentration^10^, pH^11^, enhancing superresolution^12^ and multiplexed imaging^13^.

For over a decade, the scientific community has primarily relied on Globals for Images SimFCS software developed by Prof. Enrico Gratton for phasor analysis^14^. However, maintenance and updates for this software were discontinued in 2021, prompting developers to create alternative phasor analysis tools. While several research laboratories have developed custom tools to address specific challenges^15–24^, and commercial brands such as Leica, PicoQuant, Becker & Hickl, FLIM Labs, and ISS have incorporated phasor analysis into their software suites, there are few reports on open-source software solutions. Existing platforms, based on Phyton (PhasorPy^15^, FLUTE^18^, Phasor Identifier^19^, FLIMPA^25^), Java (FLIMJ^20^)or MATLAB (PAM^17^), have successfully incorporated traditional phasor analysis techniques, such as intensity thresholding, image filtering, color-mapping, and cursor analysis (manual clustering). Despite these developments, recent advances in machine learning-clustering for image segmentation^26^ and unmixing of fluorescent species present in the same pixel via higher harmonics^27^, are not yet available in open-source formats.

To address this gap, our team has developed GSLab, a pioneering, open-source software platform designed to provide researchers with a comprehensive set of tools for advanced lifetime and spectral phasor image analysis. GSLab not only incorporates traditional phasor analysis techniques, but also introduces unique and advanced capabilities, including automated machine learning-based clustering in phasor space for image segmentation, real-time monitoring of both image and phasor space, and quantitative unmixing of multiple fluorescent species from a single pixel, applicable to any imaging system, whether commercial or custom-built.

GSLab is developed in MATLAB. It offers robust computational and visualization capabilities, allowing researchers to enhance their phasor analysis workflows effectively. GSLab compiles these tools into a user-friendly program, featuring a graphical user interface that simplifies complex analyses. MATLAB’s extensive library of built-in functions facilitates rapid development, enabling users to tailor GSLab to their specific research needs. While a MATLAB license by Mathworks Inc. may be a barrier for some, the widespread availability in academia ensures that most researchers can benefit from the advanced functionalities of GSLab. We have uploaded GSLab in a public repository^28^ under MIT license.

Figure 1 provides an overview of GSLab’ capabilities. It highlights its key features across three main areas: Input/Output, Basic Analysis, and Advance Analysis. The Input/Output section illustrates the software’s compatibility with various image formats for lifetime and spectral data, along with its export options for phasor plots and color-coded intensity images. The Basic Analysis section illustrates the commonly available tools such as cursor analysis, phasor filtering and image manipulation, enabling users to interact with and process the data efficiently. The current open-source solutions allow for the implementation of a subset of these analysis tools. The Advanced Analysis section emphasizes the software’s powerful functions for unmixing multiple fluorescence components, machine-learning based clustering, and real-time examination of the reciprocity between image and phasor space, demonstrating its robust analytical capabilities. A more detailed description of GSLab’ capabilities is available in the Supplementary Material.

**Figure 1.**
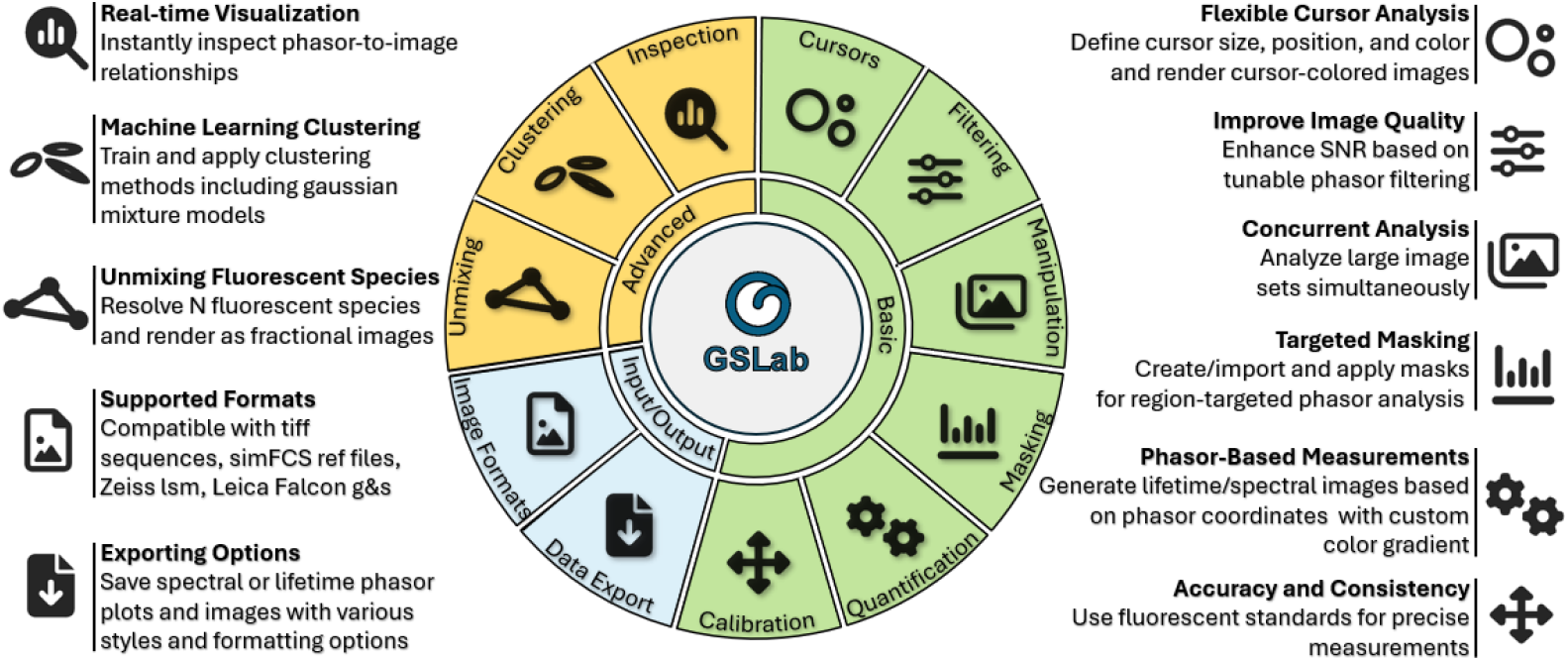
GSLab capability overview. 1) **Input/Output:** Supports major image formats for lifetime and spectral data. Allows exporting of phasor plots and color-coded intensity images based on phasor coordinates, component unmixing, or clustering results, with various styling options for both phasor plots and images. 2) **Basic Analysis**: Offers cursor analysis, phasor filtering, image manipulation, and the ability to create or load masks. Users can measure pixel phasor coordinates and create color gradients on the phasor space, generate lifetime and spectral images based on the color gradients, and calibrate the data using reference files. 3) **Advanced Analysis**: Enables unmixing of multiple fluorescence components, machine learning-based clustering of phasor distributions, and real time inspection of the reciprocity between image and phasor space.

Figure 2 illustrates the advanced capabilities of phasor analysis for automated machine learning, clustering, and unmixing of fluorescent components. We demonstrate these capabilities by analyzing two sets of data. First, label-free FLIM images of a fresh human skin specimen (discarded tissue from surgery), acquired with a custom-built clinical multiphoton microscopy device^29^ are used to demonstrate automated clustering and image segmentation (Figure 2 A-D). Second, time-resolved fluorescence images of labeled cell cultures, obtained using another custom-built FLIM-based multiphoton microscope for thick tissue imaging^30^ are used to demonstrate unmixing of fluorescent components (Figure 2 E-H).

**Figure 2.**
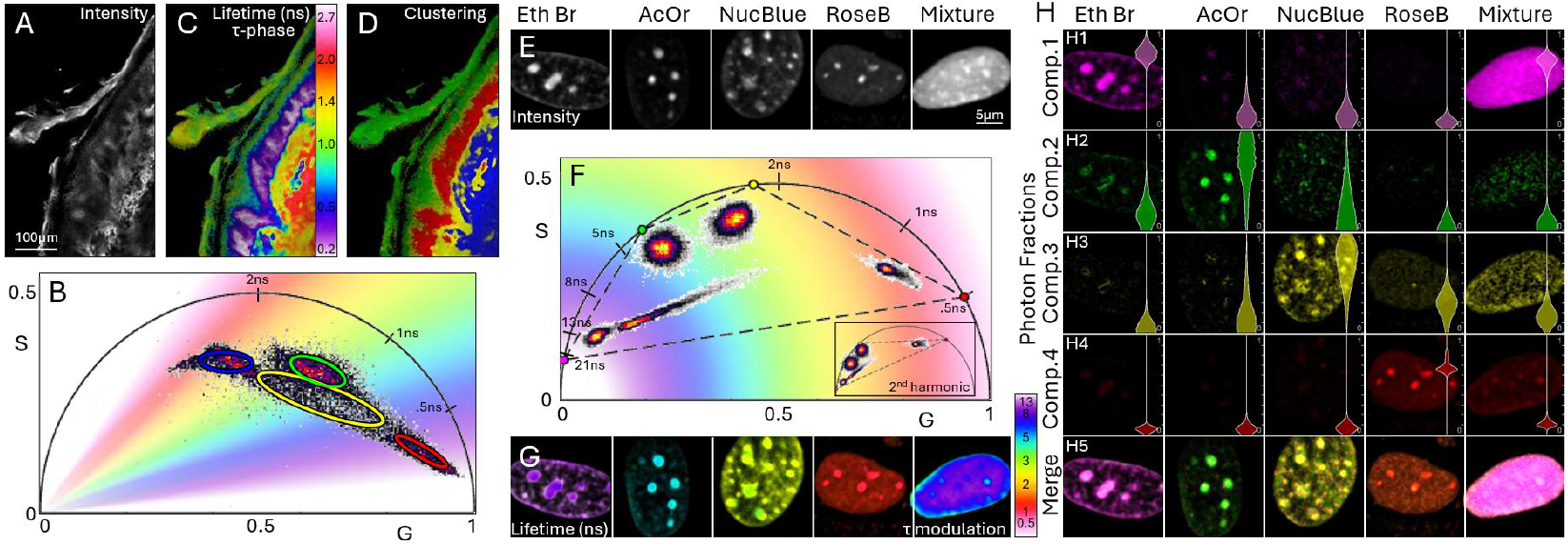
Advanced phasor analysis: Automated machine-learning clustering of phasor distributions and fluorescent component unmixing. **A**) Intensity obtained through time-resolved two-photon excited fluorescence signal detection from a human skin specimen. **B**) Corresponding fluorescence lifetime phasor plot with gradient color distribution representing tau-phase values. The plot demonstrates automated machine learning clustering into four populations with ellipses outlining 88% of pixels within each Gaussian component. **C**) The intensity image from (A) shown as color-coded lifetime image, based on tau-phase values from the phasor plot. **D**) The intensity image from (A) displayed as color-coded image highlighting the four populations identified through automatic clustering. These populations correspond to known skin structures: keratin in epidermal keratinocytes (green), melanin in pigmented keratinocytes (red), elastin in the dermis (blue), and other structures characterized by a mixed fluorescence lifetime distribution, representing a linear combination of the other three clusters. **E**) Intensity images obtained through time-resolved two-photon excitation signal detection from five cell cultures: four of them stained independently with one of the nuclear dyes Ethidium Bromide, Acridine Orange, NucBlue, and Rose Bengal, and the fifth with a mixture of all four dyes. **F**) Corresponding fluorescence lifetime phasor plot with gradient color distribution representing tau-modulation values. The plot shows the fluorescence lifetime signatures for each dye and the signature of the mixed sample as an elongated distribution in the center. The theoretical lifetime of the four dyes is used to compute the coordinates for each of the four components, depicted by the vertices of a quadrilateral. Inset displays the same data in the 2nd harmonic required for the fluorescent component unmixing. **G**) Intensity images from (E) shown as color-coded lifetime images, based on tau-modulation values from the phasor plot. **H)** Results of fluorescent component unmixing: **H1-H4**) Unmixed photon fraction images for each component (rows) and each measurement (columns). Inset violin plots illustrate pixel value distribution for each image. **H5**) Merged images for each measurement using linear addition of panels (**H1-H4**).

Phasor analysis of the skin specimen’s FLIM image (Figure 2 B) shows automated machine-learning-based clustering into four populations, which are subsequently illustrated in the corresponding color-coded FLIM image revealing well-known skin structures. Phasor analysis of the fluorescence images of stained cell cultures highlights the ability to unmix four fluorescent components by clearly resolving the fluorescence lifetime signatures of the dyes and computing the pixel fractions of the components in the mixed sample. These examples demonstrate the powerful advanced capabilities of GSLab for resolving complex fluorescence signatures within a single sample.

## Online Methods / Supplementary Material

### Phasor Analysis for Fluorescence Lifetime and Hyperspectral Microscopy

In fluorescence lifetime microscopy and spectral imaging, each pixel within the image represents a measured photon distribution, either spectral or temporal, that is determined by the fluorescent molecules present at that location. In fluorescence lifetime imaging, the characteristic times the fluorophores spend in their excited states prior to transitioning back to the ground state define the shape of this curve. In spectral imaging, the wavelength associated with the energy differences between these transitions determine the curve’s profile. The phasor approach is a technique that transforms the photon distribution curve from each pixel into a point in a two-dimensional phasor space using the first two terms of the Fourier discrete decomposition^31–33^. The two dimensions of the phasor space are usually labeled S and G, and are obtained for each pixel using the following expressions, for lifetime imaging:

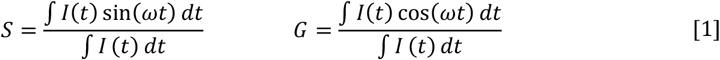

where the integrals are taken over the effective pixel dwell time *t, I(t)* is the accumulated distribution of the number of photons as a function of arrival time with respect to the excitation time, *ω=n2πT*^*-1*^ is the angular frequency such that the trigonometric functions fit *n* whole periods in the excitation period *T*, with *n* being the harmonic number of the phasor transform. Analogously for spectral imaging:

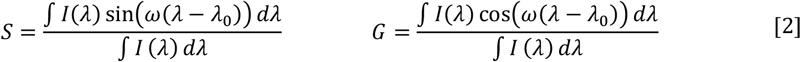

where the integrals are taken over the detection spectral bandwidth [*λ*_*0*_ *λ*_*1*_], *I(λ)* is the photon distribution as a function of the spectral dimension, *ω=n2π(λ*_*1*_*-λ*_*0*_)^*-1*^ is the angular frequency such that the trigonometric functions fit *n* whole periods in the spectral band [*λ*_*0*_ *λ*_*1*_], with *n* being the harmonic number of the phasor transform.

In practice, we operate with digital measurements; both temporal and spectral dimensions are divided into bins and each pixel is assigned a photon count for each bin. Both spectral and lifetime phasor transforms are expressed as summations over the number of bins:

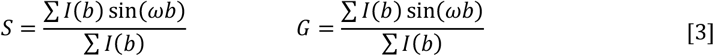

with the summations being taken over all bins with *I(b)* representing the count values at each bin *b*, and *ω=n2πB*^*-1*^ denoting the angular frequency such that the trigonometric functions fit *n* whole periods across the *B* bins.

In FLIM, if we model the intensity as an exponential decay 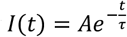, we now have a definition for lifetime: the characteristic time τ that molecules spend in the excited state. By inserting this exponential model into the equations in (1), if we solve the integrals and join the equations, we obtain the expression for the universal circle *S*^2^ = *G* − *G*^2^. Lifetime measurements on the phasor plot only make sense on this circle, any pixel that is not on the circle has a combination of lifetimes. When generating so-called lifetime images, one needs to project the phasor coordinates on to the universal circle. Using polar coordinates, we can define the phase *φ* and modulation *M* of a measurement based on the cartesian phasor coordinates:

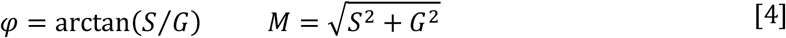

A measurement by phase is to assign the lifetime of the exponential decay such that it has the same phase as the measurement. A measurement by modulation corresponds to assigning the lifetime of the exponential decay such that it has the same modulation as the measurement. Geometrically speaking, the first is to find the point on the universal circle that has the same phase as the measurement, the second is to find the point on the universal circle that has the same modulation as the measurement (see Figure S1). We name these two lifetime measurements tau-phase and tau-modulation. The first is more precise when measuring fast lifetimes relative to the excitation frequency, the latter is more precise when measuring slow lifetimes relative to the excitation frequency. For this reason, a third method is sometimes used, namely tau-normalized^34^, which consists in projecting from the center of the universal circle. Given the phasor coordinates of a measurement *(G*,*S)*, by imposing the afore-mentioned geometrical considerations, one can derive the expressions for the three different projections, namely tau-phase, tau-modulation and tau-normalized:

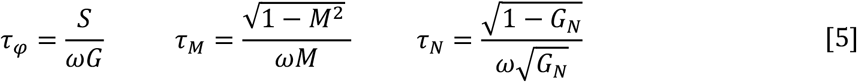

with ω = 2π*fn* being the angular frequency of the excitation source with frequency *f* (in Hertz to obtain lifetimes in seconds), and *n* the harmonic number of the transform. The parameter 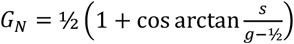 corresponds to the G coordinate of the normalized projection (see Figure S1), with the inverse tangent function returning values in the first and second quadrant.

**Figure S1.**
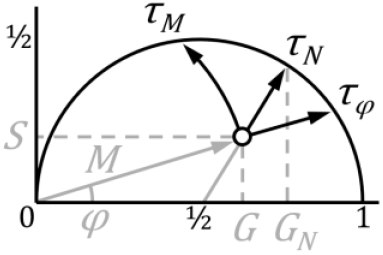
Lifetime of a measurement. Geometrical interpretation of tau -phase, -modulation and -normalized

### GSLab Capabilities

#### Input/Output

##### Image Formats

We developed our software for image analysis of lifetime and spectral measurements. We initially developed it for our home-built microscope (FLAME)^35^ operated by ScanImage (MBF Bioscience). However, we have since expanded its capabilities to support other types of image file formats for FLIM data generated by commercial microscopes such as Leica-Falcon LAS X (S and G phasor exports) and for home-built microscopes using SimFCS. Additionally, we designed the software to load and analyze hyperspectral images obtained with Zeiss ZEN (.lsm).

##### Data Export

GSLab has flexibility to export phasor plots, phasor-colored images and spreadsheets of computed values in different formats. Intensity images can be rendered using a logarithmic intensity scale. A pseudo-high dynamic range routine is implemented to compute the local dynamic range for each pixel, enabling the export of images with an extended dynamic range. Raw grayscale images can be exported or color-coded in various ways: based on cursor positions, phasor coordinates, or the results of clustering or unmixing algorithms. Colors can be applied to the image space using two methods: hard masking, which assigns a solid color to each pixel and soft masking, which creates an intensity-weighted color image. Additionally, built-in options for exporting phasor plots are available, such as the ability to adjust the binning resolution of phasor space histogram and choose between the linear or logarithmic phasor plot representation. Phasor distributions can be normalized to each loaded file to represent multiple files at the same time. Users can select different colormaps to depict the phasor density representations and for phasor measurement images. Contour lines can also be added to the phasor plots.

#### Basic Analysis

##### Calibration

To allow for an appropriate calibration and referencing of phasor coordinates, GSLab allows loading calibration samples (measurement of a fluorophore solution of known single-exponential lifetime). The user can associate each calibration file with a file name tag to identify physical detectors and/or spectral channels in the instrument. We have incorporated the use of ref file format, allowing us to save image phasor coordinates in multiple harmonics, which adds compatibility with legacy SimFCS.

##### Quantification

Phase and modulation are classical parameters for characterizing fluorescence lifetime in the frequency domain^36^. They have a direct representation on the phasor plot and are used as a way to render lifetime images. Researchers report tau-phase or tau-modulation images or a combination of both, which correspond to the lifetime associated with a measurement based on phase difference or modulation factor of the fluorescence signal with respect to the excitation signal (see Figure S1). We implement a color-coded gradient representation of these parameters, allowing users to assign each pixel a specific lifetime value and export color-coded images depending on the assigned lifetime values. The software also allows the use of custom projections on the universal circle, by setting the coordinates of the origin and selecting if projecting either radially or angularly, which would be a generalization for lifetime measurements.

##### Image manipulation

Inspired by SimFCS, our software can perform multi-image phasor analysis, enabling users to load any number of images and display them in one panel while viewing their corresponding phasor distribution in another, allowing for real time interaction between the image and phasor spaces. Users can select or deselect files from the set to display the phasor or to use the phasor transform data with the other available tools. Additionally, the dynamic range of the intensity image can be adjusted to enhance certain regions.

##### Image Masking

Phasor representation with image masking via thresholding involves selecting pixels in the image space based on their intensity to represent them on the phasor space. This functionality allows users to select or remove very bright populations or very dim/noisy pixels, which may be disguising relevant information. In combination with the previous methods, it allows users to create an image mask which can be used to exclude specific regions from analysis. Additionally, users can load image masks created in other software to pre-segment the images for phasor analysis.

##### Phasor Filtering

We have included various options to facilitate phasor analysis, including phasor filtering. Users can define a kernel size and the number of times the filter needs to be applied. The kernel is convolved over the phasor space coordinates to generate the density histogram. Additionally, users can set the resolution of the phasor space, along with the harmonic number of the phasor transform. The filtering is performed for all harmonics simultaneously.

##### Cursor Analysis

Traditional cursor analysis can be seen as manual clustering in the phasor space. GSLab implements this by allowing users to point and click to create circles (cursors) of any size on the phasor space and assign colors to these cursors. The software highlights the pixels in the image space whose phasor coordinates fall within the cursors based on the assigned color of these cursors. We have implemented these capabilities inspired by SimFCS with improvements towards ease of use.

#### Advanced Analysis

##### Inspection Mode

GSLab has a real-time built in functionality that makes use of the phasor reciprocity principle^37^ for image content exploration. The reciprocity principle connects the phasor space to the image space, allowing the selection of a set of coordinates in one space to be highlighted in the other. The cursor analysis described above uses one direction of this reciprocity principle; however, it does not operate in real time. GSLab implements an inspection mode allowing users to hover the mouse over the image space to see the corresponding pixels in the phasor space in real time, and vice versa, hover the mouse over the phasor space to see where the pixels fall on the image space. Multiple images can be loaded simultaneously which may carry some computational burden. The software addresses this by down sampling the loaded images thus reducing the computation time for all other functionalities. When exporting data, users can select between a fast, low-resolution version using the down-sampled images or a high-resolution version based on the original dimensions of the images. Users can also adjust the level of down-sampling as needed.

##### Clustering in the Phasor Space

Gaussian mixture models are the preferred machine learning algorithm for automated clustering of populations in the phasor space^27^. Automatic selection of populations enables the segmentation of structures in the image space based on their phasor signatures, facilitating further analysis based on the segmentation. This process can be seen as an automated version of the cursor analysis described above. We have implemented the gaussian mixture model algorithm, allowing users to train a model using a set of data and then apply it to other datasets. Users can define starting points for the clusters and assign colors to export segmented images based on the probability of assignment to each cluster, as determined by the posterior of the model.

##### Phasor Unmixing

Phasor unmixing in lifetime imaging is a method that allows users to compute the photon fraction for each individual species contributing to the fluorescence in a pixel^26^. Similar to phasor analysis, phasor unmixing is model-free. The user needs to input only the lifetime values of the components for unmixing. We have implemented the computation and representation of N-harmonics, allowing users to perform phasor unmixing for up to 2N+1 components. Users define a set of pure components, assign a lifetime value to each of them and solve the unmixing at each pixel. Component fractions can be exported in spreadsheets and unmixed images can be generated, both as independent fractions and as component color-coded images. Additionally, component ratio-metric images can be generated. Furthermore, users can take empirical measurements and select them as pure components, which is particularly useful for unmixing spectral components.

## Acknowledgements

We wish to recognize the impactful career of Prof. Enrico Gratton, whose contributions have significantly influenced our work. We also acknowledge the funding support for this project from the NIBIB (R01EB026705), NIAMS (R21AR082648) and NCI (R01CA259019), as well as the Skin Biology Resource-Based Center at the University of California, Irvine (P30AR075047).

## Author Contributions

A.V. and S.R conceived the project. A.V. designed and wrote the code for GSLab. A.V., B.T. and S.R. troubleshot the successive implementations. B.T., A.F.D., A.D. and N.R. performed the test experiments. A.V. and M.B wrote the manuscript with the input of all authors. M.B. supervised the project.

## Ethics Declarations

MB is a coauthor of a patent owned by the University of California, Irvine (UCI) related to the development of clinical multiphoton microscopy technology. Additionally, MB is a cofounder of Infraderm, LLC, a startup spin-off from UCI focused on commercializing clinical multiphoton microscopy imaging platforms that may benefit from the use of advanced analysis tools. The Institutional Review Board and Conflict of Interest Office of UCI have reviewed patent disclosures and found no concerns. The other authors declare no competing interests.

